# Multi-modal communication: Song sparrows increase signal redundancy in noise

**DOI:** 10.1101/750307

**Authors:** Çağlar Akçay, Michael D. Beecher

## Abstract

Although the effects of anthropogenic noise on animal communication have been studied widely, most research on the effect of noise in communication has been on communication in a single modality. Consequently, how multimodal communication is affected by anthropogenic noise is relatively poorly understood. Here we ask whether song sparrows (*Melospiza melodia*) show evidence of plasticity in response to noise in two aggressive signals in acoustic and visual modalities. We test two hypotheses: (1) that song sparrows will shift signaling effort to the visual modality (the multi-modal shift hypothesis), and (2) that they will increase redundancy of their multi-modal signaling (the back-up hypothesis). We presented male song sparrows with song playback and a taxidermic mount with or without a low-frequency acoustic noise from a nearby speaker. We found that males did not switch their signaling effort to visual modality (i.e., wing waves) in response to the noise. However, the correlation between warbled soft songs and wing waves increased in the noise treatment, i.e. signals became more redundant. These results suggest that when faced with anthropogenic noise, song sparrows can increase redundancy of their multi-modal signals, which may aid in robustness of the communication system.

## Background

Animal communication often involves signals in multiple sensory modalities [1-3]. These multi-component, multi-modal signals allow signalers to enhance information transmission compared to unimodal signals [4]. For instance, signal components in one modality may modulate the information carried in another modality (e.g., the McGurk effect in which the perception of a vowel is modified by the articulatory gesture [5]). Multi-modal signals may also be redundant, when the signal components in different modalities carry the same information (this is the standard use of the term redundant in animal communication [6, 7], which differs somewhat from the use of the term in systems biology, see [8] for a discussion of the terms). When signals have components in multiple modalities with the same meaning, communication may be more robust in noisy conditions [9].

Communication under noisy conditions has been a major focus of research recently because of concerns with the effects of anthropogenic noise on animal behavior. Noise generated by human activities can drown out animal signals [10, 11], change the characteristics of signals [12, 13] and impact fitness negatively as a result [14-16]. Most of the research on the effect of noise on signals has been focused on signals in a single modality; how noise affects multimodal signals is not well studied [17].

Multi-modal signaling may offer animals some flexibility in dealing with noise. If animals have multi-modal and redundant signals, they may shift their signaling from the noisy modality to the other, less noisy modality [7], and receivers may pay more attention to the less noisy modality [18]. There are only a few studies however that test whether such multi-modal shifts occur in nature and whether they are based on plasticity in signaling behaviors [19].

In the present study, we examine multi-modal signaling of aggressive intent in song sparrows (*Melospiza melodia*). Males signal their aggressive intent with two signals: low amplitude “soft” songs and rapid fluttering of the wings called “wing waves”. Soft songs (particularly warbled soft songs) and wing waves are often given together as part of a multi-modal display called “puff-sing-wave” [20] such that the rates of the two displays are correlated across individuals. The rate at which wing waves and soft songs are given both predict a subsequent attack on a taxidermic mount [21-23], indicating that they are redundant.

There are two types of soft songs: warbled and crystallized soft songs. Crystallized soft songs are simply regular broadcast songs sung at a low amplitude, while warbled soft songs are not found in the regular broadcast song repertoire, have a distinct syntax and notes not commonly found in crystallized soft songs, and are sung at lower amplitude than crystallized soft songs, never at broadcast amplitudes [24] (see Figure 1). Although the two different types are often given in the same stream of acoustic signaling, warbled soft songs appear to be the effective element signaling aggressive intent: Males respond more aggressively to warbled soft songs than to loud songs [25, 26], but do not respond more aggressively to crystallized soft songs than to loud songs [27].

**Figure 1.**
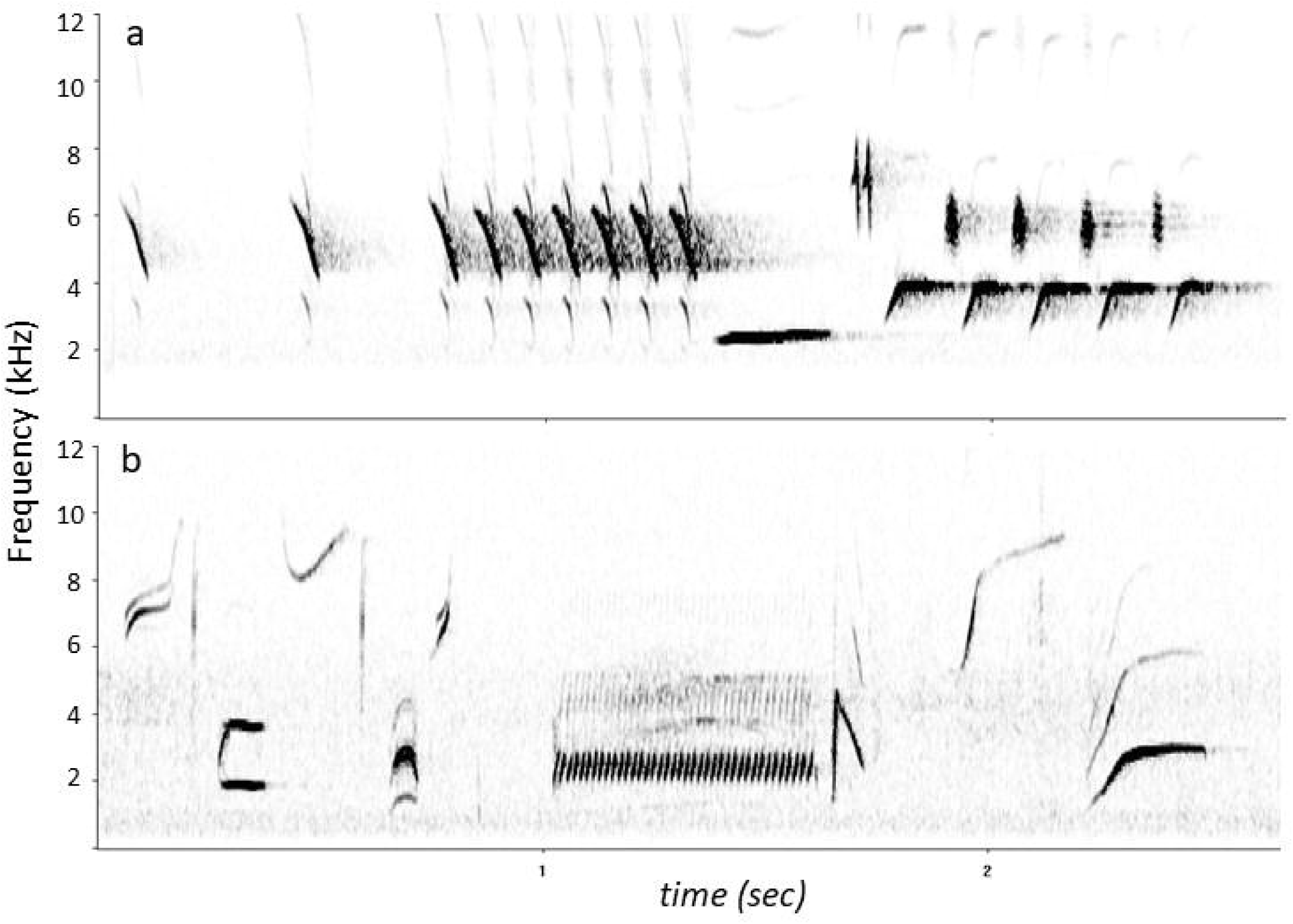
Example of a) a broadcast repertoire song of a song sparrow, and b) warbled soft song. The crystallized soft songs are structurally identical to broadcast songs but are sung at a lower amplitude. Note the alternating high and low frequency note and the broader frequency range in the warbled soft song.

We test two hypotheses on the use of multi-modal signaling under noise during agonistic interactions with a simulated intruder. The ***multi-modal shift hypothesis*** [7] predicts that when confronted with an intruder in noisy acoustic conditions, males should increase their signaling effort in the visual modality (i.e., increase rates of wing waves) since the visual signals would not be affected by acoustic noise. Simultaneously they are expected to decrease acoustic signaling via soft songs. The ***back-up signal hypothesis***, predicts that when faced with noise in one modality signalers should increase signal redundancy to maximize transmission of information [9]. Under this hypothesis we expect a stronger correlation between the visual and acoustic signals in experimentally increased acoustic noise.

## Materials and Methods

We tested 24 male song sparrows holding territories in Discovery Park, Seattle, in June 2019. The sample size was determined by an *a priori* power analysis with G*Power [28] to detect an effect size of d=0.6 with β=0.8 based on our previous research demonstrating plasticity in wing wave rates [26, 29]. We tested each subject twice about 24 hrs apart, once with noise and once without noise in a counter-balanced order.

We prepared the noise stimulus by first generating a white-noise file with Audacity software and then filtering it with the amplitude spectrum of vehicle noise recorded at our site (Marantz PMD660 and Sennheiser ME67/K6 shotgun microphone) using the package seewave [30]; see supplementary material for relative amplitude spectrum and the noise file. Stimulus songs were songs from stranger males recorded at least five years prior (all stimulus birds had disappeared by 2019). Each subject received the same song in both trials. Each stimulus song was used on one subject (except one that was mistakenly used twice).

Before each trial we measured the ambient noise according to the methods described in [31] with VLIKE VL6708 Sound level meter (A weighting, fast response). Ambient noise was below 50 dB in all trials. We then staged simulated intrusions via playback of song and a taxidermic mount either with playback of acoustic noise or without. A Bluetooth speaker (VicTsing Inc.) and the mount were taped to a natural perch about 1.5 m off the ground in the territory of the subject at least 15 m away from any boundary. We played the song stimulus from this speaker at a rate of 1 song every 10 seconds and at a maximum amplitude of 85 dB SPL, measured at 1 m with the same equipment as above. The noise stimulus was played from a second speaker (Pignose Model No. 7-100R) placed on the ground directly below the mount, face up, at 75 dB SPL measured at 1 m. These levels create an ambient noise range between 60 to 70 dB SPL within 5m of the mount which corresponds to noise levels on territories near highways [32]. We placed this speaker for both noise and control trials, but only turned it on in noise trials. The noise playback was started once the subject was within 5m of the speaker (all subjects approached to 5 m within 1 minute of their first response) and lasted until the end of the trial including the post-playback period or attack.

During the trial we recorded the behavior of the bird (Marantz PMD660 and Sennheiser ME67/K6). Two observers noted wing waves, loud songs, soft songs, and distance to the mount with each flight. Loud and soft determination for soft song was made in the field by an experienced observer (CA) which produces a reliable cut-off point [24], although the amplitude variation in song sparrow song is continuous. Playback lasted for three minutes or until the subject attacked. If the subject did not attack during playback, we waited for another 2 minutes with no song stimulus (two subjects attacked during this period). The average playback duration (SD) was 147 (64) and 143 (67) seconds for control and noise conditions, respectively.

From the recordings, we extracted the proportion of time spent within 1 m of the speaker, and counts of wing waves, loud songs, warbled soft songs and crystallized soft songs. We converted the counts of behavior into rates to account for the unequal observation durations.

All analyses were carried out in R (see supplementary materials for code). To test the multi-modal shift hypothesis we carried out paired permutation tests (1000 permutations) on wing waves, warbled soft songs, crystallized soft songs and loud songs as the response variables using the package ez [33]. The pairwise slope plots and Gardner-Altman estimation plots (supplementary materials) were made using the website EstimationStats [34]. To test the back-up signal hypothesis we ran a linear mixed model with wing waves as the response variable and subject as the random effect. We entered condition (control vs. noise) and warbled soft songs (continuous) as well as their interaction as predictor variables. We ran a second model using crystallized soft songs instead of warbled soft songs. We also report repeatability of the signaling behaviors calculated with package rptR [35].

## Results

Song sparrows did not change their overall rate of wing waves or the other signals in response to experimental noise (Table 1, Figure S2). The linear mixed model showed a two-way interaction between noise treatment and rates of warbled soft songs (Table 2. Figure 2): wing waves were significantly and positively correlated with warbled soft songs in the noise treatment (Spearman ρ= 0.56, p=0.004, n=24) but not in the control treatment (Spearman ρ= 0.37, p=0.08, n=24). The linear mixed model with crystallized soft songs showed no significant main effect or interaction effect. Eight and nine subjects attacked the mount during noise and no noise trials, respectively (6 subjects attacked in both trials, 3 only in no noise trials, and 2 only in noise trials). All signaling behaviors were highly repeatable (see supplementary materials).

**Table 1.** Results of the permutation tests, and effect sizes for the main aggressive behavior and signaling variables. Positive effect sizes indicate higher mean values in the noise condition. See supplementary materials for the slope graphs.

| Response variable | p-value (1000 permutations) | Effect size (Hedges' g) [95% CI] |
| --- | --- | --- |
| Proportion of time within 1m | 0.307 | -0.247 [-0.856, 0.3] |
| Wing waves | 0.328 | -0.126 [-0.774, 0.377] |
| Warbled soft songs | 0.118 | -0.294 [-0.853, 0.27] |
| Crystallized soft songs | 0.445 | 0.148 [-0.373, 0.76] |
| Loud song | 0.574 | 0.0885 [-0.541, 0.613] |

**Table 2.** Results of the linear mixed models with wing waves as the response variable. The predictor variables were condition (noise vs. control), warbled (left side) or crystallized soft songs (right side) and the interaction of condition with either warbled or crystallized soft songs.

|  | warbled soft songs |  |  | crystallized soft songs |  |  |
| --- | --- | --- | --- | --- | --- | --- |
|  | Coefficient (SE) | *t* | *p* | Coefficient (SE) | *t* | *p* |
| soft song | 0.46 (0.32) | 1.41 | 0.173 | 0.16 (0.45) | 0.36 | 0.73 |
| Condition (Noise) | -1.74 (1.06) | -1.63 | 0.12 | -0.87 (1.22) | -0.71 | 0.48 |
| Soft song*Condition | **0.85 (0.39)** | **2.18** | **0.04** | 0.00 (0.45) | 0.00 | 0.99 |

**Figure 2.**
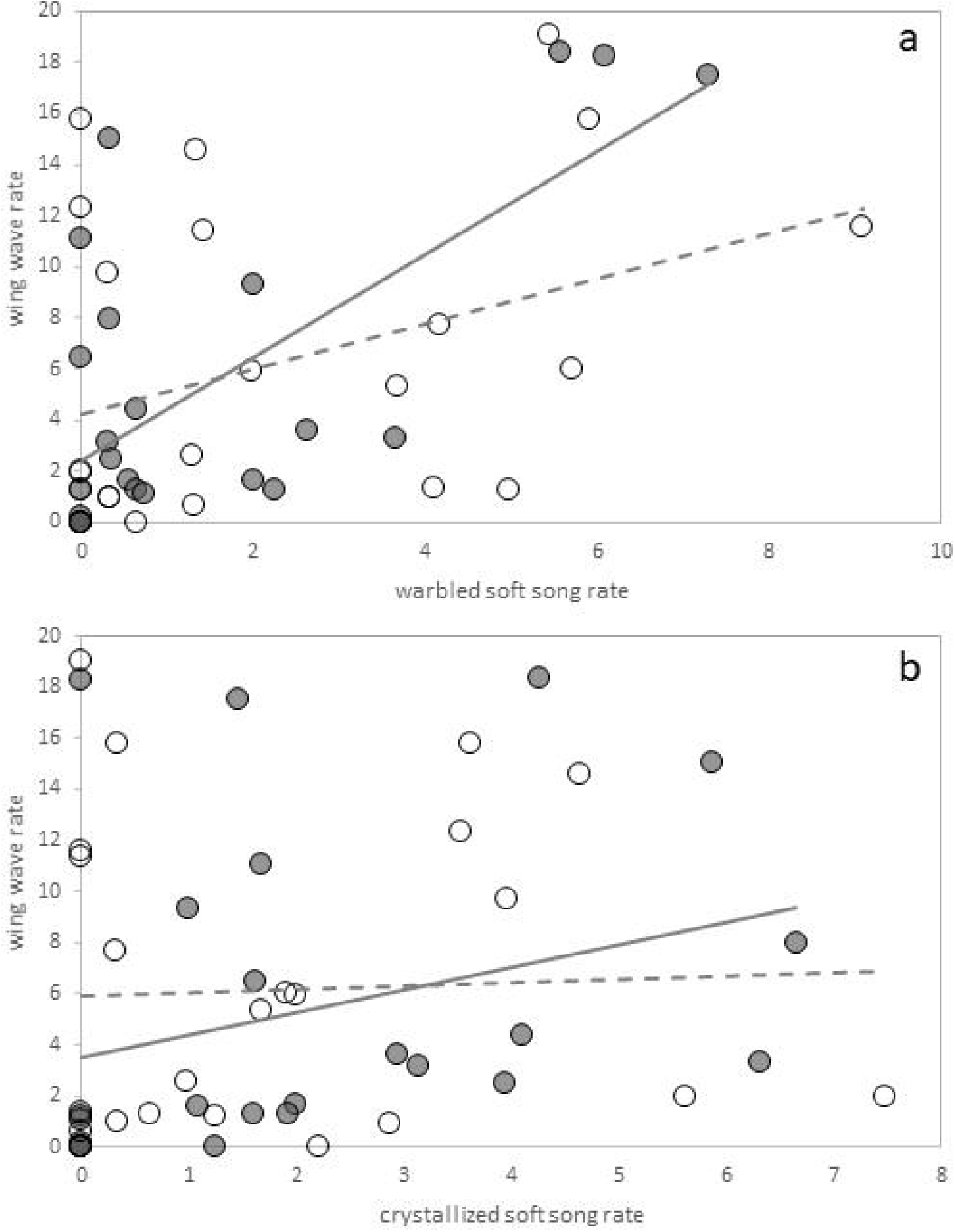
Relationship between wing waves and a) warbled soft songs, b) crystallized soft songs. The interaction of condition and soft songs is only significant for warbled soft songs. Closed circles and line is noise condition, open circles and dashed line is for control condition.

## Discussion

Song sparrows did not increase the rates of wing waves, their visual signal, when faced with an intruder in acoustically noisy conditions, contrary to the expectation of the multi-modal shift hypothesis. Interestingly a recent study comparing signaling in urban and rural song sparrows found evidence of a potential multi-modal shift: urban song sparrows gave proportionally more wing waves than rural song sparrows in response to an intruder [23]. The present failure to find a multi-modal shift suggests that the multi-modal shift in urban song sparrows is not due to short-term plasticity in response to noise, although it is possible that long-term exposure to noise may still cause a multi-modal shift.

We found that the relationship between the rates of wing waves and warbled soft songs became stronger in experimentally administered noise compared to when there was no noise. This is consistent with the back-up signal hypothesis given that two signals of equivalent meaning but in different modalities become more tightly correlated with each other in noise, ensuring redundancy in information transfer. The effect was only present for warbled soft songs, and not crystallized soft songs. This may be because warbled soft songs appear to be the effective signal of aggressive intent, with males specifically responding more aggressively to it [25, 26]. In addition, warbled soft songs tend to have the lowest amplitude (<60 dB SPL at 1m [24]), which may make them more prone to interference from acoustic noise. For these reasons coupling warbled soft songs with wing waves more tightly in noise may enhance the robustness of aggressive signaling for males. It is also possible, albeit speculative at this point, that coupling warbled soft songs more tightly with wing waves under noise may also help to enhance the reception of warbled soft songs by drawing the attention of the receivers to the auditory signal [36, 37].

Although we examined the changes in signaling effort in response to noise, we do not know how the receivers’ responses to wing waves and soft songs may change in noisy conditions. Receivers may start paying more attention to the visual signals even in the absence of an overall increase in signaling effort. In a recent aquarium study with the painted goby (*Pomatoschistus pictus*), visual signaling by males did not increase with added noise (and in fact both acoustic and visual signals were decreased in noise). Females however, appeared to pay more attention to the visual courtship in the noise condition, suggesting that visual signals are potentially more effective under acoustic noise [18, 38]. Furthermore, as in our study, there was a similar increase in the correlation between acoustic and visual signals under noise.

Further studies are needed to assess receiver responses to visual signals either coupled with or dissociated from acoustic signals under different levels of noise. Such a study could be done with a robotic model [39]. The two hypotheses (back-up hypothesis and attentional enhancement hypothesis) can also be contrasted by manipulating the temporal coordination of the acoustic and visual signals that the robotic model gives.

## Conclusion

In summary, under naturalistic field conditions, male song sparrows responded to experimentally increased noise during a simulated intrusion by increasing the correlation between their acoustic and visual aggressive signals. These results are consistent with the back-up hypothesis and suggest that a visual signal (wing waves) may provide cross-modal attentional enhancement of an auditory signal (warbled soft songs). We agree with Halfwerk and Slabbekoorn [17] that future studies on the effect of noise in signaling should consider how noise in different modalities affect signaler and receiver behavior when signaling occurs in more than one modality.

## Supporting information

Analysis code in R to reproduce results

## Acknowledgements

We thank Eliot Brenowitz for loaning us a sound level meter calibrator.

